# Sensory expectations shape neural population dynamics in motor circuits

**DOI:** 10.1101/2024.12.22.629295

**Authors:** Jonathan A. Michaels, Mehrdad Kashefi, Jack Zheng, Olivier Codol, Jeffrey Weiler, Rhonda Kersten, Jonathan C. Lau, Paul L. Gribble, Jörn Diedrichsen, J. Andrew Pruszynski

## Abstract

The neural basis of movement preparation has been extensively studied during self-initiated actions where motor cortical activity during preparation shows a lawful relationship to the parameters of the subsequent action^1,2^. However, movements are regularly triggered and constantly corrected based on sensory inputs caused by disturbances to the body or environment. Since such disturbances are often predictable and since preparing for disturbances would make movements better, we hypothesized that expectations about sensory inputs also influence preparatory activity in motor circuits. Here we show that when humans and monkeys are probabilistically cued about the direction of a future mechanical perturbation, they incorporate sensory expectations into their movement preparation and improve their corrective responses. Using high-density neural recordings, we establish that sensory expectations are widespread across the brain, including the motor cortical areas involved in preparing self-initiated actions. The geometry of these preparatory signals in the neural population state is simple, scaling directly with the probability of each perturbation direction. After perturbation onset, a condition-independent perturbation signal shifts the neural state leading to rapid responses that initially reflect sensory expectations. Based on neural networks coupled to a biomechanical model of the arm^3^, we show that this neural geometry emerges through training, but only when the incoming sensory information indicating perturbation direction coincides with – or is preceded by – a condition-independent signal indicating that a perturbation has occurred. Thus, motor circuit dynamics are shaped by future sensory inputs, providing clear empirical support for the idea that movement is governed by the sophisticated manipulation of sensory feedback^4^.

## Main

Humans and animals are often able to prepare a movement in advance and such preparation generally makes movements better. The neural basis of movement preparation and its relationship to movement execution has frequently been studied with delayed action paradigms, where the nature of a future movement is instructed but its execution must wait until a subsequent go cue (for review^1,2,5^). During the preparatory period, between the movement instruction and the go cue, muscle activity remains unchanged but motor cortical activity represents parameters of the future movement^6–15^, predicts movement variability^16^ and reaction time^17–20^, and is causally linked to motor execution^5,21,22^, presumably by setting the initial state of the dynamical system that ultimately produces movement^10,23–27^.

Although preparing specific movement parameters is an essential aspect of self-initiated actions, movements are regularly triggered or corrected based on sensory inputs caused by disturbances to the body or environment. Since such disturbances can often be predicted based on experience or by other streams of sensory input, and since preparing for potential disturbances would improve motor performance, we hypothesized that sensory expectations should also directly shape preparatory activity in motor cortical areas. Such a scheme is a key prediction of modern theories of biological motor control based on optimal feedback control^1,4,28^ and would be consistent with previous reports that motor cortical areas rapidly respond to sensory inputs (for review^29,30^) in a way that accounts for biomechanical^31^ and task constraints^32–37^.

Here we show that, when cued about the likely direction of future mechanical disturbances, humans and macaque monkeys readily incorporate expectations about the upcoming sensory input into their movement preparation and that this preparation improves their performance. We then demonstrate that information about sensory expectations is widespread in monkey motor circuits and that the geometry of these sensory expectation signals relates to the neural dynamics that ultimately produce corrective responses to mechanical disturbances. Lastly, we develop a normative model of the motor system showing precisely how this neural geometry is beneficial for countering disturbances and how it relies on the timing of incoming sensory signals.

### Long-latency stretch reflexes are sensitive to sensory expectations

To investigate whether expected sensory inputs shape preparatory activity in motor cortical areas, we designed a task where human participants were given probabilistic information about how a future mechanical disturbance would move their arm relative to a goal target. We specifically chose mechanical disturbances that evoked stretch reflexes because the various components of the stretch reflex involve different neural circuits^38,39^. The short-latency component, measured as muscle activity occurring 20-50 ms after muscle stretch, is generated entirely by spinal circuits. The long-latency component, measured as muscle activity occurring 50-100 ms after muscle stretch, includes a contribution from motor cortical areas via the transcortical feedback pathway^29,40^. Thus, if motor cortical circuits are set in accordance with sensory expectations, we should find that the long-latency stretch reflex is sensitive to sensory expectations. Indeed, our approach builds directly on previous studies showing that long-latency stretch reflexes are shaped by many of the same parameters known to influence preparatory activity in motor cortical areas in the context of self-initiated movements^31–34^.

Human participants (N=20) performed the task in a KINARM exoskeleton robot (Methods, Fig. 1a). They maintained the position of their hand within a small central target and were given an explicit visual cue about the probability that their elbow joint would be flexed or extended following a variable delay period (Fig. 1b). Following this variable delay, a perturbation drawn randomly from the cued probability distribution was applied and participants had to respond to the perturbation by moving their hand into the goal target as quickly and accurately as possible.

**Fig. 1.**
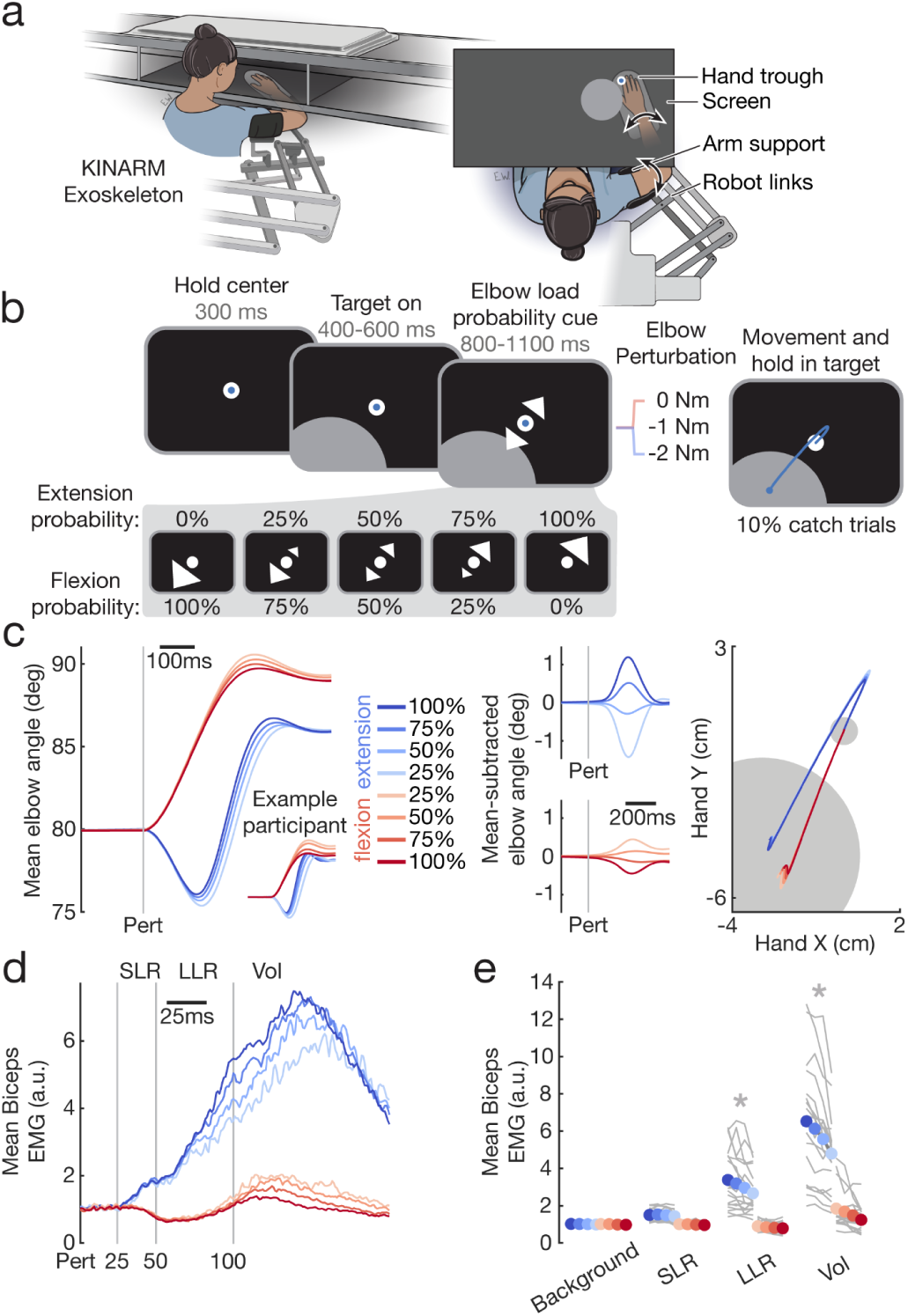
| Long-latency stretch reflexes take into account sensory expectations. a, Human participants were seated in the KINARM exoskeleton robot, allowing presentation of visual stimuli in the horizontal plane, tracking of the arm, and application of mechanical forces to the shoulder and elbow joints independently. **b**, Participants were required to hold their hand at a small central target and were shown a peripheral goal target. On single trials participants then received one of five visual cues indicating the probability that the upcoming mechanical perturbation would flex or extend their elbow joint and thus push their hand into or out of the goal target. After a variable delay, a perturbation drawn from the displayed probability distribution was applied and subjects had to respond to the perturbation by moving their hand into the goal target quickly and accurately. **c**, Mean elbow kinematics across participants (N=20) show that participants responded to the perturbations in a graded fashion, where the speed that subjects moved to the target depended on the cued probability of each perturbation. **d**, Mean biceps muscle activity measured via surface electromyography (EMG) as a function of perturbation direction and probability cue. Note that the initial excitatory response to extension perturbations in the short latency reflex (SLR) window is the same for all probability cues, but that the subsequent excitatory response in the long latency reflex (LLR) and Voluntary (Vol) windows is scaled by the probability cues. **e**, Mean muscle responses were significantly modulated by probability cues in the LLR and Vol windows (repeated-measures ANOVA, p < 0.001).

Participants were very good at the task (success rate: 85.7 ± 6.3%; mean ± s.d.), and rarely initiated a movement incorrectly during catch trials without perturbations (error rate: 5.0 ± 3.5%). The probability cue biased task performance in a graded fashion, with participants responding to perturbations faster as the probability cue indicated a greater certainty of a particular perturbation direction (Fig. 1c). Mechanical disturbances evoked excitatory or inhibitory stretch reflex responses starting ∼20 ms after muscle stretch depending on whether the disturbance lengthened or shortened the muscle (Fig. 1d, Fig. S1). Notably, the evoked responses were initially insensitive to probability information, which appeared to emerge in a graded fashion ∼70 ms after muscle stretch. A repeated measures ANOVA comparing average muscle activity in pre-defined epochs as a function of probability confirmed this observation (Fig. 1e). We found no reliable effect of probability in the short-latency epoch (SLR, 20-50 ms after muscle stretch; F(3,19) = 2.35, p = 0.08). In contrast, we did find a reliable effect of probability in both the long-latency (LLR, 50-100 ms; F(3,19) = 7.55, p = 0.0002) and voluntary (100-150 ms; F(3,19) = 18.28, p <0.0001) epochs. Note that this modulation did not appear to reflect anticipatory modulation of muscle activity prior to the disturbance^41^, as there was no reliable effect of probability on muscle activity just prior to muscle stretch (background, -200-0 ms; F(3,19) = 0.86, p = 0.46). Could participants have been making binary guesses about the most likely perturbation on individual trials, causing the average to show a graded response? This possibility is highly unlikely, as the distribution of time elapsed to reach the target following perturbation onset did not significantly differ from a unimodal distribution for any participant or probability (Hartigan’s dip test, all p > 0.1).

### Preparatory neural states in motor cortical areas represent perturbation probability

Our finding that long-latency stretch reflexes are sensitive to sensory expectations strongly suggests motor cortical involvement but cannot establish a direct link to its preparatory state. To directly test whether sensory expectations shape preparatory activity in motor cortical areas, we trained two macaque monkeys in an almost identical version of the task performed by human participants using a monkey version of the KINARM exoskeleton robot (Methods, Fig. 2a). Unlike human participants, who were verbally informed about the probabilistic nature of the cues and enacted the association immediately, monkeys learned to associate the visual cues with probabilities through experience over several months.

**Fig. 2.**
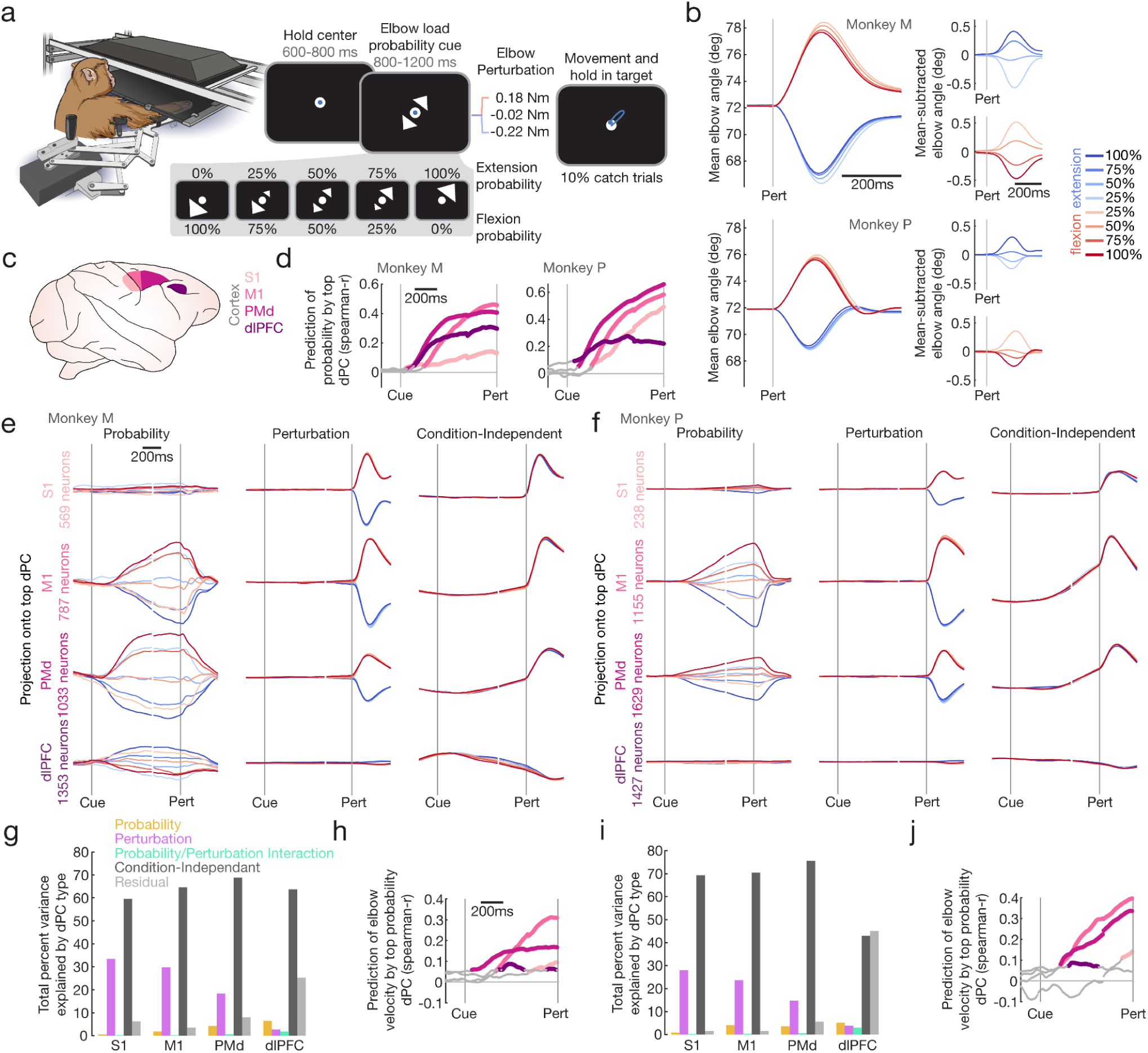
| Preparatory neural states in motor cortical areas represent perturbation probability. **A**, Two macaque monkeys performed a version of the task very similar to human participants using a non-human primate version of the KINARM exoskeleton robot. **b**, Mean elbow kinematics across recording sessions showed clear scaling by probability cue in both perturbation directions. **c**, Single neuron recordings were made over sessions using Neuropixels probes from four cortical areas, primary somatosensory cortex (S1), primary motor cortex (M1), dorsal premotor cortex (PMd), and the dorsolateral prefrontal cortex (dlPFC). **d**, Demixed principal components analysis (dPCA) was performed across the single neurons of each area to disentangle task-related signals at the population level. Projections onto the top probability dPC were used to predict probability magnitudes on single trials pooled across all sessions of each area. Colored lines indicate correlations significantly greater than chance, where chance level was obtained by randomly shuffling probability conditions and correlating with the true probability condition (permutation test, p < 0.001, 10000 iterations). **e**, Top dPCs are plotted for each task factor (probability, perturbation, condition-independent) across areas for Monkey M, and **f**, Monkey P. dPCs are normalized to the max value of each task factor pooled across areas. **g**, Summary of total variance explained per task factor in each area for Monkey M. **h**, Mean prediction of single-trial elbow velocity in the 150 ms after the perturbation by the top probability dPC, performed separately for each perturbation direction, chance level calculation as in d. **i,j,** The same analyses as g,h for Monkey P.

After training, the monkeys performed the task well, completing a large percentage of initiated trials correctly (Monkey M: 89.1 ± 4.1%, Monkey P: 93.2 ± 1.9%; mean ± s.d. over sessions) and with only a small percentage of trials being aborted due to unwanted movement before perturbation onset (Monkey M: 7.3 ± 3.1%, Monkey P: 5.4 ± 1.6%). Similar to human participants, monkeys showed a clear effect of the probability cue on their elbow kinematics in the 400 ms following the perturbation (Fig. 2b, one-way ANOVA of single-trial elbow angle over probability conditions, Monkey M: extension F(3, 16424) = 79.6, p < 1e-6, flexion F(3, 16274) = 169.3, p < 1e-6, Monkey P: extension F(3, 7099) = 61.8, p < 1e-6, flexion F(3, 7054) = 35.5, p < 1e-6), but no effect of probability in the 100 ms before the perturbation (Monkey M: extension F(3, 16424) = 0.6, p = 0.60, flexion F(3, 16274) = 1.8, p = 0.14, Monkey P: extension F(3, 7099) = 0.2, p = 0.92, flexion F(3, 7054) = 0.7, p = 0.55), demonstrating that they learned to prepare for future perturbations differently depending on the probability cue. As with human participants, it’s unlikely that these differences were caused by guessing what perturbation would occur on individual trials, since the distribution of time elapsed to reach the target during elbow perturbations did not significantly differ from a unimodal distribution (Hartigan’s dip test, each condition and session tested separately – Monkey M: all p-values > 0.1, Monkey P: all p-values > 0.05).

Having established that monkeys use these probabilistic cues to prepare their responses to perturbations, we recorded single neurons from multiple brain areas to assess how neural population coding supported this preparation. To do so we developed a new recording setup allowing high-throughput single neuron extracellular recordings using Neuropixels probes (see Methods). In both monkeys, we recorded from four areas potentially involved in this processing (Fig. 2c): the primary somatosensory cortex (S1), primary motor cortex (M1), the dorsal premotor cortex (PMd), and the dorsolateral prefrontal cortex (dlPFC), recording from 8191 single neurons in total across areas and monkeys. To decompose the signals in the neural population related to the task, we applied demixed principal components analysis (dPCA) to the firing rates of populations of neurons in each brain region and monkey separately. dPCA allows us to separate dimensions related to probability information (the cues presented to the monkey), the perturbations delivered, condition-independent changes, and any linear interactions. Figure 2d shows the single-trial correlation between the top probability dimension of each area and the perturbation probability throughout the preparation time. Between the cue and perturbation time, the top probability dimension was predictive of probability to some extent in each area, however, there were clear differences across areas. Notably, the first areas to show a single-trial correlation with probability were PMd and dlPFC, around 100ms after the cue was presented, followed by a ramping of probability information in M1. On the other hand, representations in S1 were small and ramped up latest during preparation.

To visualize the nature and extent of these representations, we plotted the condition-averaged neural population activity of each area projected into the top dPC of each factor (Fig. 2e-f). During preparation conditions corresponding to identical visual cues overlap in the probability dimension, leading to 5 distinct traces. The representations of probability information were clearest in PMd and M1, reaching their peak shortly before the perturbation and then decreasing throughout the movement period. As expected, the probability dimensions represented only probability information and did not delineate what perturbation was actually applied, demonstrating that probability information was linearly separable from perturbation information. Even though dPCA simply tried to find the neuronal dimension that maximally distinguished between the probability cues, the emerging dimension ordered these cues clearly by the probability magnitude. Thus anywhere where there were differences between cues, the neural representation scaled directly with the probability of each perturbation direction, showing a straightforward geometry of activity that related conditions to each other by probability magnitude. The perturbation dimension clearly separated the two perturbation directions in all areas except in dlPFC, where representation of perturbation direction was very small. Also notable is that all areas showed dominant condition-independent signals. These varied across areas, with S1 and M1 showing the sharpest responses to the perturbation while higher order areas (PMd and dlPFC) showed less sharp responses to the perturbation and greater ramping during preparation.

The origin of the condition-independent signal at perturbation onset is unclear (see Discussion), since the majority of sensory inputs from the periphery would be directional and show an opposite response for different elbow perturbation directions.

To summarize these results across all dPCs, we looked at the total variance explained by each factor (including interaction and residual variance) across areas and monkeys (Fig. 2g,i). We found a clear gradient of probability information across areas, with the least proportion of variance explained by probability in S1, with more explained variance in M1, PMd, and then dlPFC. On the other hand, we saw the reverse gradient for perturbation information (most dominant in S1, least dominant in dlPFC). The interaction of probability and perturbation explained only a very small amount of variance overall. Condition-independent dimensions represented the majority of variance explained in all areas, highlighting the importance of these time-varying signals and in line with previous work during self-initiated movements^42,43^. An analysis of perturbation detection time showed the earliest condition-independent responses were in S1 and M1 (Fig. S2, 15.5 ms and 18 ms post-perturbation, respectively). Residual variance not captured by dPCA was overall very small with the exception of dlPFC, suggesting that the remaining variance is either not linearly separable or related to other factors not experimentally controlled.

Finally, if the probability dimensions extracted with dPCA causally relate to the magnitude of the behavioral response to the perturbation, we would expect that as probability representations fluctuate on single trials, these would have an effect on behavior on the same trials. To test this, we used the single-trial projections of neural activity onto the first probability dimension and correlated them with elbow velocity in the first 150 ms following the perturbation period (Fig. 2h,j, correlations were done separately for each perturbation direction and averaged). In previous analyses, probability information was present in dlPFC and S1. However, we did not observe any clear correlation between single-trial fluctuations in probability information and subsequent elbow velocity after the perturbation. On the other hand, neural activity close to the perturbation in both M1 and PMd significantly predicted elbow velocity after the perturbation on single trials, suggesting that the probability representation in these areas directly modulates rapid muscle responses to perturbations, a result consistent with their key role in the transcortical feedback pathway.

### Preparatory neural states in motor but not sensory thalamus represent perturbation probability

Given the ubiquity of the sensory expectation signal in multiple cortical areas we tested whether this signal is already present at a subcortical level. To gain some insight into this question, we extended our recording setup to allow the use of 4.5 cm Neuropixels^44^ in Monkey P. Thalamus is particularly interesting given the rich and differentiated input and output connectivity of its various nuclei. We targeted the ventroposterior lateral thalamus (VPL), the ventral lateral posterodorsal thalamus (VLpd), and the ventral lateral anterior thalamus (VLa). VPL receives tactile and proprioceptive information from the cuneate nucleus and has dense projections to S1^45^ and M1^46^, while VLpd receives primarily cerebellar input and projects to the the supplementary motor area (SMA) and PMd/M1^47^, and VLa receives pallidal input and projects to SMA/PMd/M1^47,48^, motivating us to record from SMA as well. Surprisingly, probability information was present in almost all of these areas rapidly after cue presentation (Fig. 3b). Specifically, VLa and VLpd showed an increase in probability prediction in the top probability dPC within 100ms of the cue, followed closely by SMA. On the other hand, the VPL, the sensory nucleus, showed only low levels of prediction much later in preparation, similar to what we observed in S1.

**Fig. 3.**
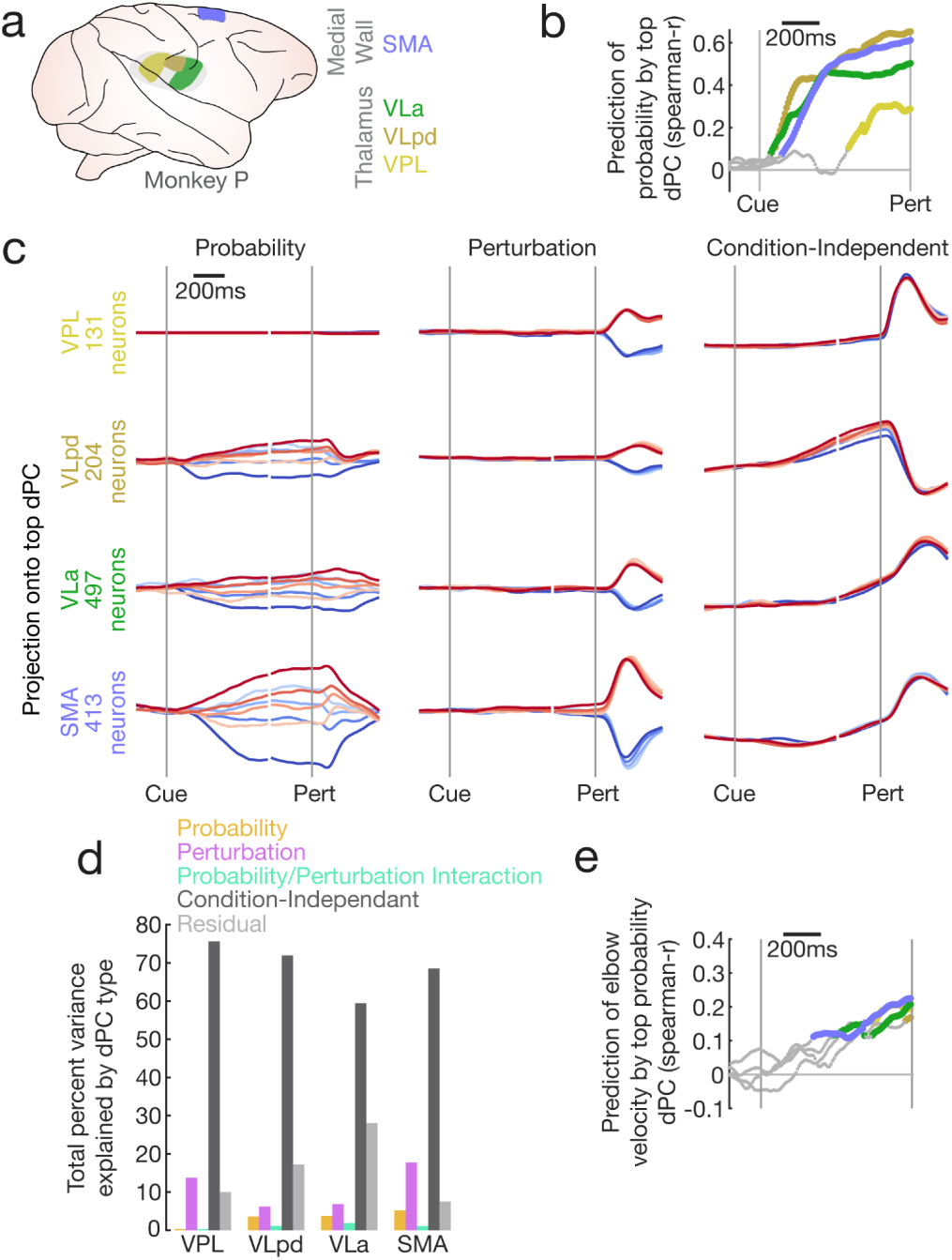
| Preparatory neural states in motor but not sensory thalamus represent perturbation probability. a, In addition to previous recordings, single neuron recordings were made from Monkey P over sessions using Neuropixels probes from four areas, the supplementary motor area (SMA) in the medial wall, and the VLa, VLpd, and VPL nuclei of the thalamus. **b**, Demixed principal components analysis (dPCA) was performed across the single neurons of each area to disentangle task-related signals at the population level. Projections onto the top probability dPC were used to predict probability magnitudes on single trials pooled across all sessions of each area. Colored lines represent significant correlations, where chance level obtained by randomly shuffling probability conditions and correlating with the true probability condition (permutation test, p < 0.001, 10000 iterations). **c,** Top dPCs are plotted for each task factor (probability, perturbation, condition-independent) across areas. dPCs are normalized to the max value of each task factor pooled across areas. **d**, Summary of total variance explained per task factor in each area for Monkey P. **e**, Mean prediction of single-trial elbow velocity in the 150 ms after the perturbation by the top probability dPC, performed separately for each perturbation direction, chance level calculation as in b.

A summary of variance explained (Fig. 3d) shows that, as in previous analyses, condition-independent information was by far the most dominant, and the timing of these responses clearly showed that the perturbation was detectable earliest in VPL (Fig. S2, 15 ms post-perturbation). SMA and VPL had the largest perturbation responses, and probability information in VPL accounted for less than 0.3% of total variance.

Visualizing the projections in the top dPC for probability (Fig. 3c) confirms that probability information was not dominant in the VPL. The geometry of probability information in the other thalamic nuclei and SMA were similar to previous results and organized by relative probability. Interestingly, post-perturbation elbow velocity was weakly predictable from the top probability dPC of all these areas close to perturbation time (Fig. 3e), suggesting some direct dependence between neural state and subsequent behaviour.

### Probability representations are widespread during preparation and are replaced by prediction error and perturbation direction representations during movement

In almost all areas investigated the neural population data were organized such that the 100% probability conditions of opposite perturbation directions were the farthest away from each other in neural space and the other probability conditions were positioned in a graded fashion in between these extremes (Fig 4a, Probability Model), indicating a direct representation of relative probability. However, predictive coding^49,50^ theories predict that some components of the neural response would represent how incongruous perturbations are with expectations (Fig. 4a, Prediction Error Model), scaling directionally based on how much the delivered perturbation deviated from expectation. Similarly, these theories predict the presence of a signal representing how surprising a given perturbation was (Fig. 4a, Unsigned Prediction Error Model), as both of these prediction error-related models have been shown to be relevant for learning and memory^50^. Responses could also have differed purely based on actual perturbation direction (Fig. 4a, Perturbation Model). To test the relative contribution of these models to explaining neural data, we converted all models and neural population data into euclidean distance matrices between all pairs of conditions (relational dissimilarity matrices, RDMs) to represent the geometry of each model and the neural population. Disentangling the contribution of each of these models to observed neural activity is challenging due to multicollinearity that exists between models. To overcome this, we used a concept from cooperative game theory to estimate the contribution of each model. We used non-negative linear least-squares regression to predict the RDM of the neural population at each time point based on linear combinations of our model RDMs. Importantly, we exhaustively fit every possible combination of models, allowing us to calculate Shapley values, which estimate the true contribution of each model to explaining neural data (see Methods).

**Fig. 4.**
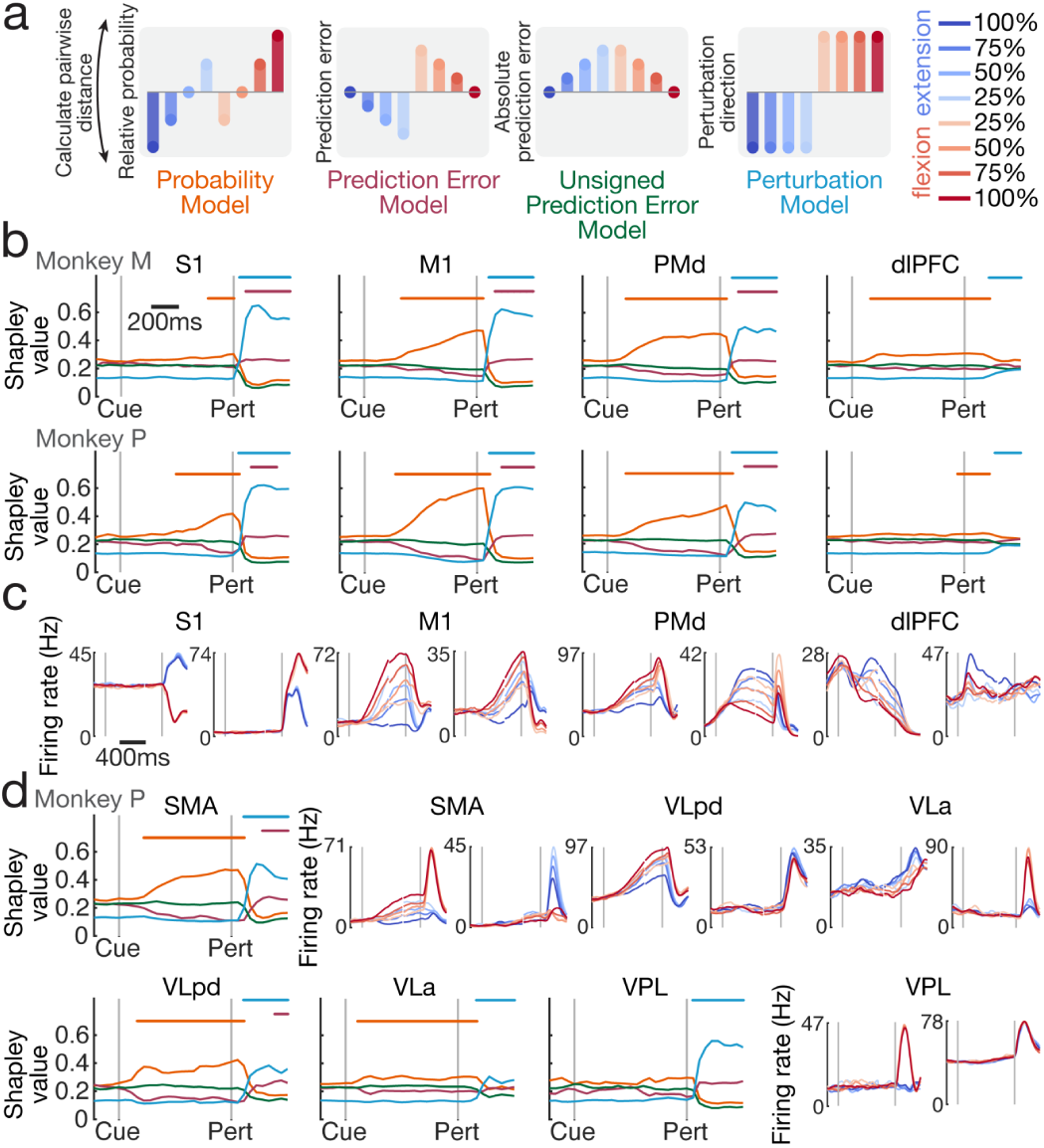
| Probability representations are widespread during preparation and are replaced by prediction error and perturbation direction representations during movement. a, We compared the ability of four potential models to explain neural population geometry by predicting the euclidean relational dissimilarity matrices (RDMs) between all pairs of conditions in the neural population space as a linear combination of model RDMs using non-negative linear least-squares regression. Neural RDMs were calculated in the high-dimensional neural space, while model RDMs were calculated in the 1-dimensional space of each model. **b,** Each model’s unique contribution to explaining neural geometry was estimated using Shapley values. Fits were considered significant (solid bars above plots) if they exceeded the 99th percentile of the null distribution generated by randomly shuffling conditions for each neuron independently and repeating the Shapley value calculation 100 times. **c,** Example mean firing rates for single neurons across areas. **d**, Model fitting results and example neurons for medial wall and thalamic areas in Monkey P.

During the preparatory period only the Probability Model was able to significantly fit the data, and this effect was widespread across cortical areas (Fig. 4b). The earliest and strongest representation was in PMd, while the latest and weakest was in S1. Probability representations collapsed dramatically within the 50-100 ms after the perturbation, and in many areas were replaced immediately by the Perturbation Model and the Prediction Error Model. Although perturbation direction representations were far more dominant, prediction errors were also reliably and significantly present, and can be seen in some example single neurons (Fig. 4c, e.g. 4th neuron from the left). Perhaps surprisingly (no pun intended), in no case did the Unsigned Prediction Error Model ever significantly explain neural geometry, indicating that responses directly related to surprise were not present. In Monkey P, we performed the same analysis for additional medial wall and thalamic areas (Fig. 4d). Significant linear representations of probability were also widespread during preparation in these areas, with the exception of VPL. Most of these areas also showed a significant representation of the Perturbation Model and the Prediction Error Model, while no area showed a significant representation of the Unsigned Prediction Error Model.

### Only some areas represent sensory expectations accumulated from experience

In our previous experiments, humans and monkeys were shown a random probability cue on each trial, making the visual cue the only source of information for forming sensory expectations. Although examples like this exist in natural environments, expectations about future sensory inputs can also be discovered through experience interacting with the environment. Therefore, we designed an additional experiment to investigate the neural representation of sensory expectations acquired over multiple trials. In this experiment (Fig. 5a, Methods), monkeys received alternating blocks of randomly cued probabilities (presented visually, as in previous experiment) and adaptation blocks where perturbations were drawn from a single randomly chosen probability distribution that remained fixed for the duration of that block and was not visually cued to the monkey (*e.g.*, 75% extension). We did not include the 100% probability conditions in the adaptation blocks to maintain some uncertainty about the underlying distribution. The only way to determine the underlying perturbation probability distribution in adaptation blocks was by experiencing a succession of perturbation trials. In Monkey M, we recorded 333 neurons in S1, 399 in M1, 244 in PMd, and 1329 in dlPFC. In Monkey P, we recorded 215 neurons in S1, 586 in M1, 1629 in PMd, 1427 in dlPFC, 131 in VPL, 204 in VLpd, 497 in VLa, 413 in SMA.

**Fig. 5.**
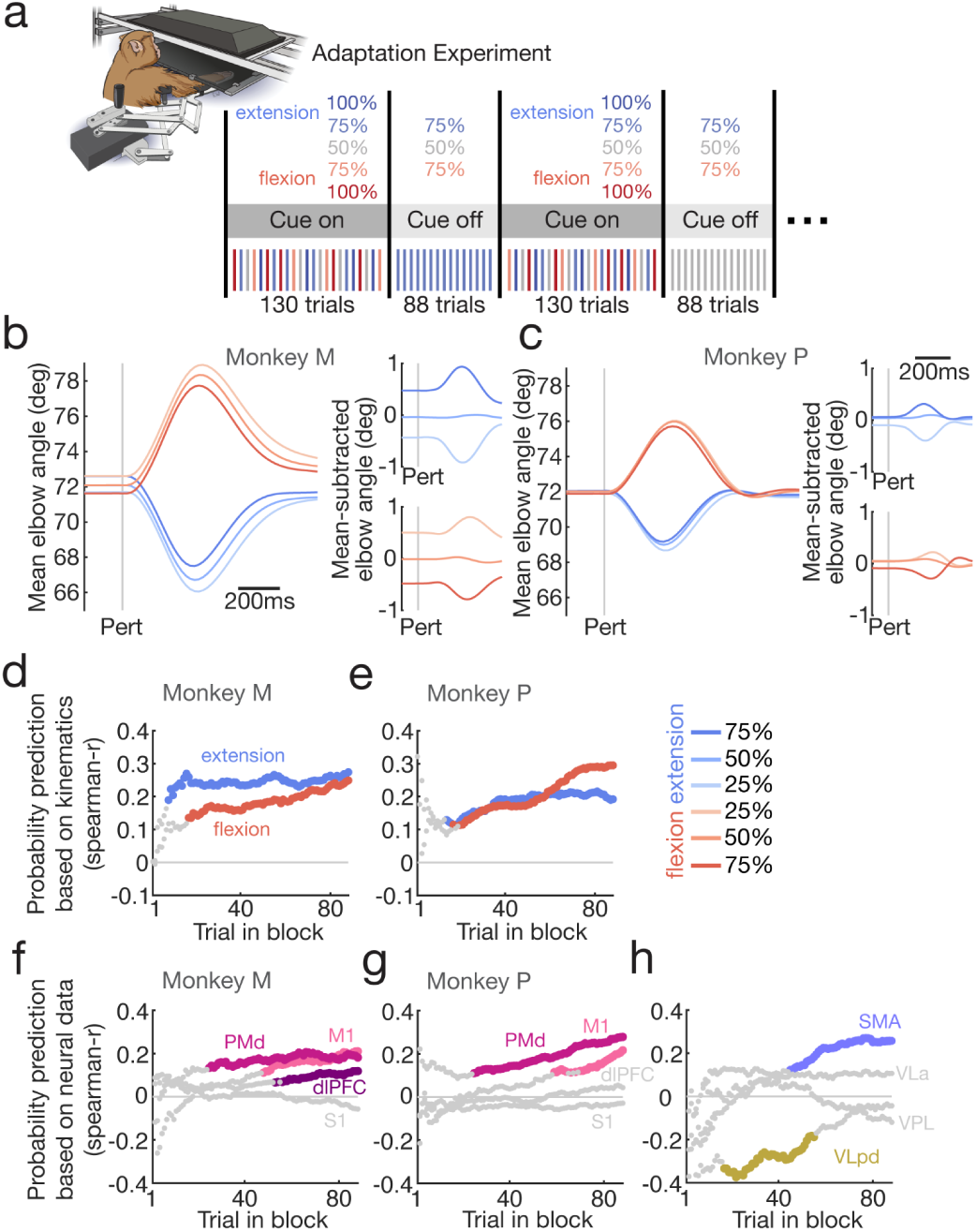
| Only some areas represent sensory expectations accumulated from experience. a, Monkeys performed an adaptation variant of the main experiment in which trials alternated in block between randomly visually cued probabilities (as in main experiment) and adaptation blocks with no probability cue in which perturbations were drawn from a randomly chosen probability value (75% extension, 50/50% or 75% flexion) that remained fixed within each block. **b,c** Mean elbow angle of each condition for trials within the adaptation blocks only. **d,** we aggregated all of the visually-cued trials across sessions (for each monkey and perturbation direction separately) and fit linear regressions to predict probability based on the post-perturbation kinematics (shoulder and elbow velocity in the window 1-400 ms post-perturbation; lasso regularization parameter for least-squares linear regression selected using bayesian optimization with 5-fold cross-validation). We used this fitted regression to predict the probability condition on adaptation trials and assessed performance as the spearman correlation for trials at all points within an adaptation block. Correlations used all trials within a sliding window including the 50 trials up to and including the current trial in block. Chance level calculations for remaining panels were made by randomly shuffling (10000 iterations) the probability values of each trial for each point within a block and calculating the spearman correlation with the true value. Correlations were considered significant (solid colors) if they exceeded the 99.9th percentile of the random distribution. **f,g,** Same analysis as previous panel, but using neural population activity in the 300 ms prior to the perturbation to predict probability condition (linear regression with L2 penalty). **h,** Same analysis for additional areas in Monkey P.

Both monkeys adapted to the distribution of experienced perturbations, scaling their responses based on the probability of the underlying distribution (Fig. 5b,c). Monkey M accumulated small offsets in elbow position during adaptation blocks, but scaling by probability was still apparent in post-perturbation elbow responses (Fig. 5b, right panels). To dissect the time course of this adaptation in a quantitative way, we aggregated all of the visually-cued trials across sessions (for each monkey and perturbation direction separately) and fit linear regressions to predict the probability distribution of each trial based on the post-perturbation kinematics (shoulder and elbow velocity in the 1-400 ms post-perturbation; L1 regularization coefficient selected using bayesian optimization with 5-fold cross-validation). We used this regression fit on the visually-cued blocks to predict probability conditions on adaptation trials and assessed performance as the Spearman correlation for trials at all points within an adaptation block (Fig. 5d,e). For both monkeys, post-perturbation kinematics started showing a significant representation of probability condition 10-20 trials into the adaptation block. An optimal Bayesian integrator would reach 95% confidence in the most likely probability distribution after ∼15 trials in the case where each of the three distributions are equally likely, as was the case here.

To test which brain areas show a similar representation of sensory expectations in the visually-cued experiment and the adaptation experiment, we repeated the previous regression analysis instead using populations of neurons in each area to predict probability, training the regression on the 300 ms before perturbation onset in the visually-cued blocks and testing on the adaptation blocks (Fig. 5f,g). Here, across monkeys we found that probability representations were found only consistently in PMd and M1, starting earlier in adaptation blocks in PMd, suggesting that these areas either are involved in the accumulation of evidence forming particular sensory expectations, or receive this information from other areas. Interestingly, despite some subcortical (VLa, VLpd) and cortical (SMA) areas showing probability representations during our visually-cued experiment, only SMA and VLpd showed a clear representation of probability during the adaptation experiment. Of all areas examined, VLpd showed a significant representation of probability earliest in the block and did not show a significant representation at the end of the block. The sign of the significant correlation for VLpd was also reversed, suggesting that it may play a different role during visually-cued trials and adaptation trials.

### Models of closed-loop motor control emergently develop a simple geometry of sensory expectations dependent on feedback timing

Given the strong presence of sensory expectations in motor cortical areas, we further explored why this representation arises and how it improves motor performance. To do so, we needed to create simulations that paralleled the closed-loop nature of motor control. We used our previously published open-source toolbox, MotorNet^3^, to create closed-loop models of reaching where recurrent neural networks control realistic muscles and receive delayed sensory feedback (Fig. 6a, Methods) similar to biological organisms. Thirty-two networks were trained on a random reaching task where they had to reach between random points in the workspace after receiving unpredictable perturbations delivered directly to the joints of the arm model. Paralleling the monkey’s exposure to both normal movements in their everyday experience and the experimental task (Fig. 2), network training interleaved training iterations of the random task and the experimental task. During the experimental task, the networks also received a probability cue as in the human and monkey experiments. Delayed proprioceptive feedback allowed the model to detect displacement due to mechanical perturbations applied to the limb. Importantly, mirroring what we observed empirically in early sensory areas (VPL, S1), we included a delayed condition-independent perturbation signal as part of the proprioceptive feedback that signaled when a perturbation occurred.

**Fig. 6.**
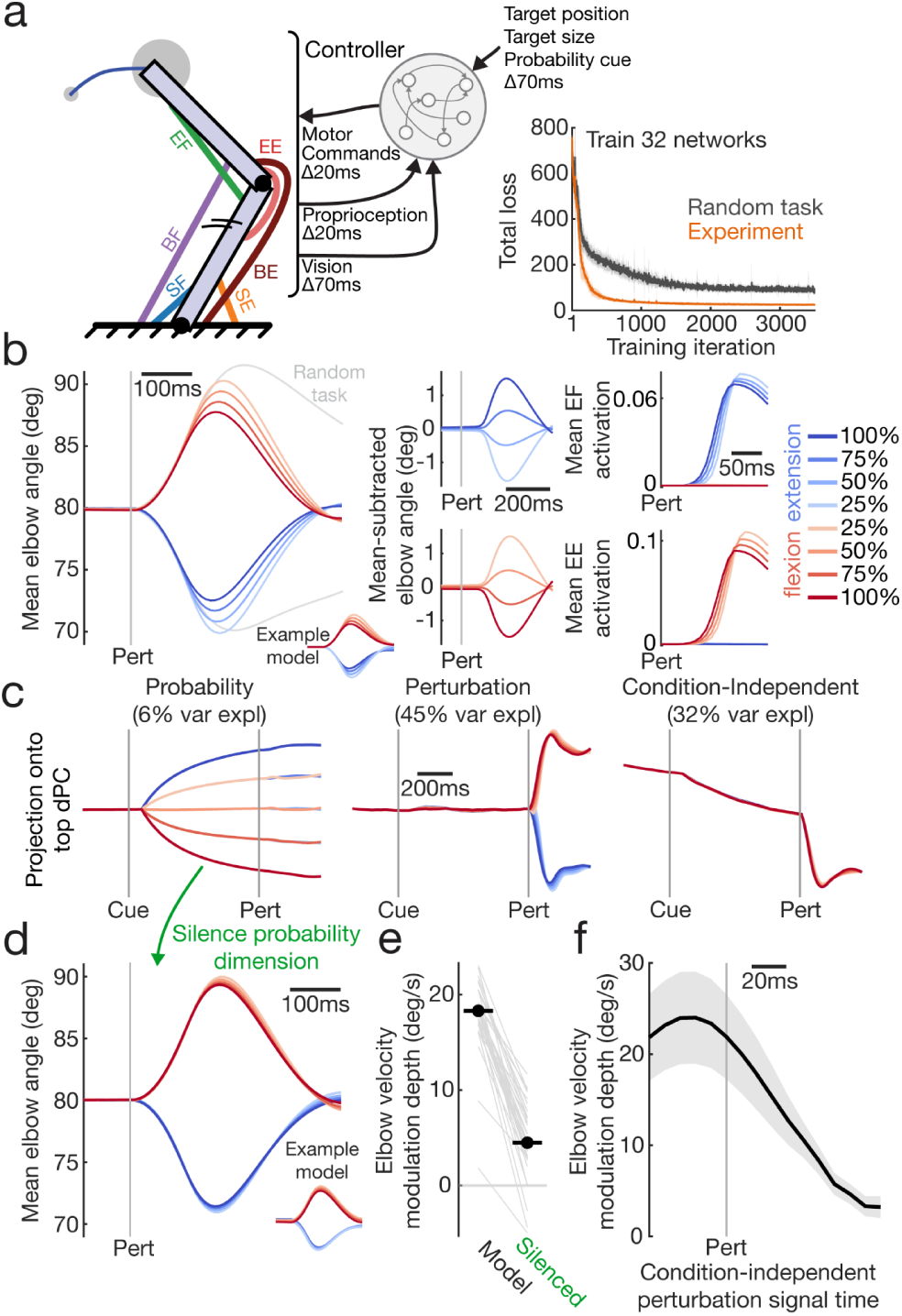
| Models of closed-loop motor control emergently develop a simple geometry of sensory expectations dependent on feedback timing. a, We used an open-source toolbox, MotorNet^3^, to train 32 recurrent neural networks to control a biomechanical model of the arm during reaching, including realistic muscles, feedback, and delays. Models were trained both on a random reaching task with targets throughout the workspace that had to be reached after a random mechanical perturbation, and a version of the experiment performed by humans and monkeys. **b,** Average kinematics of models performing the experiment scaled with probability due to scaling of muscle activation within the long-latency reflex window (50-100 ms post-perturbation). **c,** Performing dPCA on models (example model shown) revealed similar neural dimensions as observed in the empirical data. **d,** Silencing the top probability dimension during trials attenuated the ability of the networks to represent sensory expectations but did not affect the ability of the network to correct for perturbations. **e,** Quantification of the reduction of kinematic modulation depth due to sensory expectation when silencing top probability dimension. **f,** Modulating the time at which the condition-independent perturbation signal was provided to the network (original trained time was 20 ms after the perturbation) shows that as this signal arrives later, even by 10s of milliseconds, the ability of sensory expectations to affect behavior disappears, since directional perturbation information becomes available from the periphery. Shaded error represents SEM over models.

Although the models were not trained to produce a specific movement trajectory or pattern of muscle activity, the models responded more quickly to perturbations that were more likely than to perturbations that were less likely (Fig. 6b), very similar to humans and monkeys. The models learned this association between probability cues and perturbation probability by experiencing many perturbations paired with each visual cue, and similar results were obtained in networks with a one-hot input for visual cues (Fig. S3). Behavioral effects were due to scaling of muscle responses with probability quickly after perturbation onset, starting in an epoch akin to the long latency reflex window due to the sensory delays introduced into our feedback loop (Fig. 6b, right panels). Importantly, when networks were trained without the condition-independent input to inform them of perturbation onset, they were able to correct for perturbations, but they did not exhibit responses that scaled with sensory expectations (Fig. S4). The reason is that, in these networks, although perturbations were eventually detected using an internally generated condition-independent signal, they were not detected quickly enough to make use of sensory expectations before the perturbation direction was disambiguated by sensory input.

Decomposing neural activity of the model in the same way that we did for monkey data using dPCA revealed very similar dimensions for probability, perturbation, and condition-independent signals as observed in the empirical data (Fig. 6c). Importantly, the geometry of sensory expectations was simple and equivalent to the empirical data. To test if this probability dimension was causally responsible for the behavioral effects observed, we eliminated all neural activity projecting into this dimension by subtracting the appropriate amount of neural activity from each neuron at each time point in the trial, without retraining the model. Figure 6d demonstrates that when this dimension was eliminated the scaling of kinematic responses by probability was almost completely abolished. This effect was quantified by calculating the modulation depth due to probability during movement (Fig. 6e, max divergence between elbow kinematics for 100% flexion and 100% extension conditions in the -100-300 ms around perturbation onset), which confirmed that the effect of sensory expectations on motor performance were almost completely driven by this dimension.

Lastly, we wanted to understand what constraints allowed the model to take advantage of this probability information in its feedback responses. To test this, we modified the timing of the condition-independent perturbation input by manipulating its latency from its original value of 20 ms post-perturbation, without retraining the networks (Fig. 6f). The effectiveness of sensory expectations dropped very quickly as the latency increased, showing essentially no effect once it was delayed more than ∼50ms, while reducing the latency increased the effectiveness of sensory expectations. In other words, properly deploying sensory expectations during feedback responses requires that the perturbation is detected early enough to use that information before directional proprioceptive information resolves the ambiguity in perturbation direction and eliminates the need to respond based on prior expectations.

## Discussion

Our results demonstrate that humans and monkeys incorporate knowledge about future sensory inputs when preparing a movement and that this preparation improves their performance. Neural data shows that information about sensory expectations is widespread across cortical and subcortical areas, generally following a simple neural geometry that directly represents the probability of each perturbation direction. A neural network trained to control a biomechanical model of the arm reveals that incorporating sensory expectations into movement preparation is advantageous when responding to disturbances, provided that the perturbation is detected early enough to act on sensory expectations before the perturbation direction becomes clear.

How does the neural representation of sensory expectations fit into our understanding of motor cortical control^23,51^? Motor cortex has an expansive ability to represent task variables in its preparatory state^1^, including prior information about goal location^52–55^ and reward^56^, but we demonstrate for the first time that motor cortical areas directly represent expectations about sensory inputs during preparation (Fig. 7, output null trajectories). When disturbances do occur, responses are triggered by a condition-independent signal (with a similar profile to the condition-independent signal that precedes voluntary movements^43^ that quickly produces a muscle response (Fig. 7, output potent trajectories). This muscle response proportionally reflects current expectations about perturbation direction (Fig. 7, muscle activity), similar to how goal-directed movements following perturbations reflect a continually updating movement plan^57^. As in self-initiated movements, neural activity initially evolves in the neural space based on the flow field determined by recurrent dynamics. As sensory information about the actual disturbance (in our study, the perturbation direction) arrives, the modified flow field directs the neural activity onto the trajectory appropriate for the muscle activity necessary to counteract the actual perturbation. The straightforward relationship between neural preparatory state and muscle activity observed here suggests that sensory expectations should combine with goal-related preparatory states. For example, sensory expectations should interact with goal-related signals to have a strong impact on motor output when perturbations displace the limb away from a goal target, but a weak impact when perturbations displace the limb into a goal target, a prediction that can be tested directly in future experiments.

**Fig. 7.**
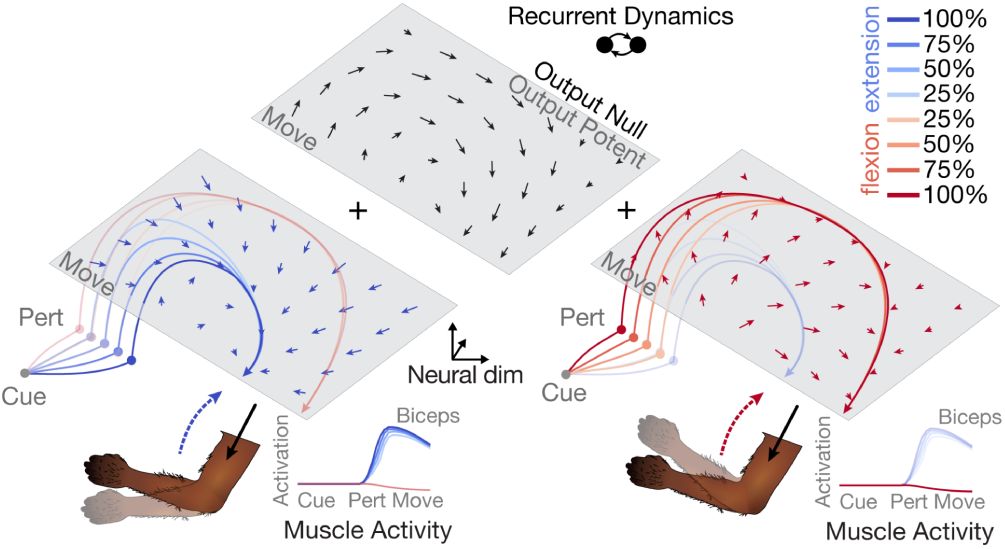
| Schematic of the neural dynamics of sensory expectations. Motor cortical areas directly represent sensory expectations during preparation as the relative probability of events (in this study perturbation direction). Once an output-potent response is triggered by a condition-independent input, muscle activity initially reflects expectations about sensory inputs as the neural state evolves through a flow field determined by recurrent dynamics. As additional sensory information about the disturbance arrives (in our case directional information), the influence of sensory feedback on the flow field pushes neural trajectories onto the path necessary to produce the muscle activity needed to counteract the actual disturbance.

In addition to the directional proprioceptive information transmitted through cortical areas following perturbations^33,34^, our results reveal a prominent condition-independent signal upon which feedback responses likely rely. What is the origin of this signal? The vast majority of information in VPL, the thalamic nucleus projecting most strongly to the primary somatosensory cortex, was condition-independent, suggesting this signal could have originated in the periphery, spinal cord, or brainstem. One possibility is that cutaneous receptors, when aggregated together at the level of second-order neurons (cuneate nucleus), could transiently signal the presence of a perturbation^58^. Pacinian corpuscles are a likely candidate due to their sensitivity to high frequency vibration and the fact that their receptive fields cover such large areas, giving them low directional resolution^59^. Another possibility is that fusimotor drive through gamma motor-neurons increases muscle spindle sensitivity such that transient vibrations during perturbations in any direction produce a condition-independent signal through Ia afferents^60^. Determining the general function of this condition independent signal and the circuit that constructs it is an important area for future work.

What implications do our results have for predictive coding, which has been proposed as a widespread mechanism for increasing sensitivity to input across the brain^61–64?^ We show that the early sensory areas investigated (VPL, S1) show very little predictive coding in our task, but very strong condition-independent signals. Predictive feedback in early sensory areas has been proposed to improve state estimation during movement in the presence of delays^65^. However, in our experiments there were no ongoing movements when perturbations arrived and therefore no corollary discharge of motor commands to sensory areas. The presence of the rapid condition-independent signal discussed above likely eliminates the need for improving state estimation immediately following unexpected perturbations, and we see no evidence that sensory predictions are used in this case to increase input sensitivity. Furthermore, the presence of prediction error signals during the response to the perturbation is likely a consequence of response speed, since the most vigorous responses will be completed earliest, leading to an increased tuning to the most unexpected perturbations later in the movement period.

Only a subset of areas showed sensory expectations when these expectations were accumulated over trials, namely areas involved with motor preparation and execution (SMA, PMd, M1) and VLpd in the thalamus, which receives primarily cerebellar input and outputs to SMA/PMd/M1^47^. The presence of probability information earliest in the cerebellar thalamus is noteworthy, as the cerebellum has been identified as a crucial component of state estimation during self-action^66–68^ and muscle responses related to the expected duration of a mechanical perturbation are eliminated in capuchin monkeys during cooling of the dentate nucleus^69^. Our results suggest that the cerebellum may be necessary for initially forming expectations in the adaptation task, but may play a different role either when probabilistic information is visually-cued or once stable sensory expectations have been formed during adaptation.

## Methods

### Human Experiment

#### Participants

Twenty healthy individuals (13 males and 7 females, aged 18-35 years, 2 left handed) participated in this experiment. All participants reported normal or corrected-to-normal vision and no history of neuromuscular impairments. Prior to data collection, all participants provided informed written consent. Participants were paid for their time and were able to withdraw from the study at any time. The study was approved by the Office of Research Ethics at the University of Western Ontario.

#### Apparatus

Participants were seated with their right arm in a KINARM robot exoskeleton (Fig. 1a, BKIN Technologies^70^), allowing flexion and extension movement of the shoulder and elbow joints in the horizontal plane. The robot can independently apply specific flexion or torques at these joints. The two segments of the exoskeleton, consisting of the upper arm and forearm, have three adjustable cuff sizes to match the dimensions of the participant’s arm. Foam pads were inserted into any remaining space between the cuffs to ensure tight coupling of the limb to the applied torques. After adjustment of the robot, calibration was performed to align a real-time, 0.5 cm diameter cursor on the right index fingertip of each participant. The hand-position feedback and visual targets of the experiment were displayed in the same horizontal plane as the arm movement. These virtual-reality images were projected in front of participants at eye-level via an LCD monitor onto a semi-silvered mirror. Before initiating the experiment, an opaque blinder was installed beneath the mirror to occlude direct vision of the physical right-arm during all trials. Kinematic data were sampled at 1000 Hz.

#### Experimental procedure

Throughout the duration of the experiment, a constant background load of 1 Nm extension torque was applied at the elbow, pre-exciting the flexor muscles. The use of a background load extending the elbow increases the stability and magnitude of flexor muscle responses^41,71^. To initiate each trial, participants moved their hand to a target (0.5cm diameter) representing the external angles of 80° and 60° for the elbow and shoulder joints, respectively. As instructed, participants tried to exert the minimum force necessary to hold their arm at the home target without co-contraction of antagonistic muscles. After 300 ms in the home target, the goal target (3.5cm diameter) appeared for a random period between 400-600 ms (Fig. 1b). The goal target was presented at a location that could be reached with a 10° pure elbow flexion from the home target. Then the arrow(s) indicating the probability of elbow perturbation direction appeared for a random period between 800-1100 ms before the perturbation was applied at the elbow. All 5 probability cues were equally likely, and randomly selected from a pool of 880 trials. The arrows were created with areas directly proportional to the percent probability. The perturbation was then applied (step-torque of ±1 Nm), which either flexed their elbow, moving their hand into the target, or extended their elbow, moving the hand away from the target. At the moment the perturbation was applied, the probability cues disappeared and visual feedback about hand location disappeared for 50ms. Participants were instructed to move to the white target once they felt the perturbations, and to do so in less than 700ms. If this was achieved, the target changed from white to green. However, if participants took more than 700ms, the target changed from white to red. This feedback was used to prompt participants to move quicker for the next trial if they moved too slowly. If participants moved off the home location prior to the perturbation, the trial was aborted and repeated later in the experiment. No restrictions were implemented on the trajectory of their arm movements. Regardless of green or red feedback, after holding their arm at the goal target for 400 ms, the torque is returned to the level corresponding to the constant background load. Participants then immediately moved to the home button to start the next trial.

Participants completed 49 practice trials, which were not included in the analysis. As part of the 880 trials, participants randomly received 10 trials of each condition as a catch trial. During catch trials, the cue appeared but the perturbation was never applied. The target automatically turned green after 2 seconds of holding the home target. Catch trials were used to ensure participants were not moving before the onset of the perturbation. Rest breaks were provided throughout each experiment at approximately 15-20 min intervals or when requested by the participants.

#### Electromyographic recording

The skin above the muscles of interest was scrubbed using a piece of gauze soaked with rubbing alcohol. The EMG electrodes (Delsys Bagnoli-8 system with DE-2.1 sensors, Boston, MA) were coated with conductive gel (Chattanooga, CA, USA, REF4248). The electrodes were taped to the skin surface above the belly of three right-arm muscles: the short head of the biceps brachii, an elbow flexor; the brachioradialis, an elbow flexor; and the medial head of the triceps brachii, an elbow extensor. The electrodes were aligned parallel to the muscle fibers. A reference electrode was secured on the left clavicle of each participant. EMG signals were amplified with a gain of 1000 and digitally sampled at 1000 Hz. The collected EMG data was then bandpass filtered at 10-500 Hz using a zero-phase, second-order Butterworth filter and full-wave rectified.

Muscle activity of elbow flexors were normalized by their mean activity from the last 200 ms prior to perturbation onset across all trials. Muscle activity of the elbow extensor, medial tricep, was normalized to mean EMG activity during three special trials at the start of the experiment. These three trials totaled 11 seconds with a constant 1 Nm elbow flexion torque.

### Non-human primate experiments

#### Subjects

Two male rhesus macaques (Monkey M, Macaca mulatta, 10 kg, 15 years old; Monkey P, Macaca mulatta, 16 kg, 11 years old) participated in the study. All procedures described below were approved by the Institutional Animal Care and Use Committee at Western University (Protocol #2022-028).

#### Experimental procedure

The design of the main monkey experiment closely mirrored the human experiment. Throughout the experiment, a constant background load of 0.02 Nm extension torque was applied at the elbow. On each trial, monkeys waited with their fingertip in a central target (located under the fingertip when the shoulder and elbow angles were 32° and 72°, respectively; target size: 1.2 cm diameter). After a variable delay (600-800 ms), one of the five possible probability cues appeared randomly. In the monkey experiment, the probability arrows were colored to further differentiate them (dark blue / 100% extension, light blue / 75% extension, white / 50% extension, light orange / 25% extension, dark orange / 0% extension). If, at any point before the perturbation, the hand went outside the home target, the trial was aborted. Trials were excluded from analysis if at any point during the delay period hand velocity exceeded 0.5cm/s. For Monkey P, these trials were aborted in real-time, while for Monkey M they were excluded from analysis. After a variable delay of 800-1200 ms, monkeys received one of two unpredictable elbow perturbations (±0.2 Nm step-torque) which served as a go cue to compensate for the perturbation and return to the central target. For monkey M, at the time of perturbation onset all visual feedback was frozen until the hand returned to the goal target. For monkey P, all visual feedback was frozen for 150 ms after the perturbation. After returning to the central target and holding the hand there for 700 ms, a liquid reward was given. In both cases the probability cues remained on until the end of the trial. In 10% of trials, after 1200 ms no perturbation was applied and a liquid reward was given. In perturbation trials, the amount of liquid given at the end of the trial scaled with the speed of the return movement. Trials in which the time between the perturbation and the reward exceeded 1.2 seconds were excluded from analysis.

#### Electrophysiological recording

We performed high-density Neuropixels probe recordings (1.0 - 1 cm, 1.0 NHP - 1 cm, and 1.0 NHP - 4.5 cm). After training on basic tasks, both monkeys were implanted with custom 3D-printed titanium implants (accurate to 0.2mm) that were designed to precisely conform to their individual skulls as determined by a model obtained using micro-CT. Titanium implants were fixed in the skull using a variable number of titanium screws and included a built-in recording chamber and head post. Neural recording targets were identified by registering the CT to a pre-surgery MRI, and identifying the 3D location of each brain area by warping segmentations from a composite macaque atlas to the individual MRI of each animal (NMT v2^72,73^, CHARM^72,74^ and SARM^75^ atlases, see Hirai & Jones^76^ for additional thalamic parcellations). The use of skull conforming titanium implants allowed us to precisely plan recording trajectories to target desired structures. The precision of our implantation technique was confirmed post-mortem in Monkey M to be accurate to within <0.5 mm on the cortical surface.

After monkeys were trained in the experiment, craniotomies were performed over the planned recording areas. In Monkey M, a large craniotomy was performed to expose the entire recording area, while in Monkey P, small 2.7 mm burr holes were drilled over recording sites as needed. In Monkey M, a custom holder was designed (Neuronitek) for use with 1.0 cm Neuropixels to allow insertion through the dura using 2-4 mm retractable guide tubes and actuated with Narishige microdrives. In Monkey P, we created a new design (Neuronitek) for use with the 4.5cm NHP Neuropixels to allow insertion through the dura using 9 mm retractable guide tubes and actuated using a manual microdrive. For each recording configuration, we 3D printed a custom holder (Formlabs 3B+, Grey resin V4) that aligned the Neuropixels along a specific, pre-defined trajectory targeting the areas of interest. Recordings in the primary somatosensory cortex (S1) primarily targeted Brodmann area 1 and area 3b.

#### Neural data processing

Neural data were recorded from Neuropixels probes using SpikeGLX. Neural data were processed using a custom processing pipeline (https://github.com/JonathanAMichaels/PixelProcessingPipeline). For Monkey M, AP stream data was first drift corrected using spike localization and decentralized registration^77,78^ implemented in spikeinterface^79^, which was able to accurately track vertical probe drift and correct it. Due to the large craniotomy, some of these recordings had large drift (0-250 um). Neural data were then processed with Kilosort 2.0^80^ to further stabilize recordings during spike sorting. For Monkey P, drift was minimal due to small craniotomies (drift 0-15 um), so we immediately processed the data using Kilosort 4.0^80^ including built-in drift correction. Single neurons were considered successfully recorded if they were flagged by Kilosort as single neurons using default parameters, and if they were stably recorded for the duration of the recording. To determine whether neurons were properly isolated over the course of the recording, we generated the average firing rates of each neuron for each condition of the main experiment (8 conditions) divided up into 5 equal blocks of trials, additionally averaging across all time in each trial (200 ms before cue onset to 300 ms after perturbation onset), which yielded a matrix of 8 x 5 values for each neuron. We then calculated the mean index of dispersion for each neuron (variance over time block divided by mean over time block, averaged across conditions) to estimate how stable each neuron was over the course of the recording. It’s important to note that this metric does not test neurons for tuning to the task, only reliable responses over the course of the recording. Neurons with an index of dispersion below 2 were included in further analysis. The majority of neurons had an index of dispersion <1, and shifting this threshold ± 1 did not affect results.

#### Demixed principal components analysis

Principal component analysis (PCA) is commonly employed to reduce the dimensionality of high dimensional datasets by finding a low dimensional representation that captures large amounts of variance using independent linear combinations of neurons. For PCA, given a matrix of data *X*, where each row contains the average firing rates of one neuron for all times and task conditions, PCA finds an encoder, *F*, and an equivalent decoder, *D*, which minimizes the loss function 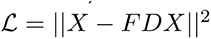 under the constraint that the principal axes are normalized and orthogonal, and therefore *D* = *F*^*T*^. Unfortunately, data that is represented in this way often heavily mixes the effect of different task parameters between latent dimensions. We would like to extract dimensions that dissociate our specific task conditions. To achieve this, demixed principal components analysis (dPCA) was performed^29^ using freely available code: http://github.com/machenslab/dPCA. In contrast to PCA, dPCA seeks to explain marginalized variance with respect to our specific task variables (probability, perturbation, and time), instead of merely explaining total variance. Unlike PCA, dPCA utilizes a separate encoder and decoder, such that the loss being optimized was 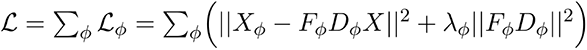, where *X*_φ_ is the marginalization of the full data with respect to each of our task parameters of interest and the λ term is a regularization parameter, preventing overfitting. Marginalizations of *X* can be obtained by averaging over all parameters which are not being investigated and subtracting all simpler marginalizations. In our case the marginalizations of interest were probability *x* time, perturbation *x* time, time, and probability *x* perturbation *x* time. The specific value of *λ* was determined using 5-fold cross-validation for each brain area in each monkey, allowing each factor to have a different value of *λ*_*φ*_.

dPCA requires data for all combinations of levels of each factor, which was not fully the case for our data since in the 100% probability conditions of each perturbation direction the opposite perturbation direction never occurred. To handle this small amount of missing data, we used a technique proposed in the original dPCA paper and fit a generalized linear model to each neuron at each time point, using the task factors (probability and perturbation) as a design matrix. In order to match trial-to-trial variability, firing rates included random Gaussian noise that scaled with the standard error of each model coefficient. While this simulated data was used for fitting dPCA, in no case was it used during analysis or calculation of variance explained.

#### Assessing relative model contributions to neural geometry

Due to the multicollinearity of the four explanatory models we used in Figure 4, we employed Shapley values derived from cooperative game theory^81,82^ to estimate the true contribution of each model. To compute the Shapley value *φ*_*i*_ for each predictor *i*, we use the formula: 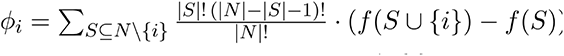, where *N* is the set of all predictors, 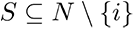 represents each possible subset of predictors that excludes *i*, 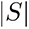 is the size of subset *S*, and *f*(*S*) is the performance metric (in our case, R-Square) achieved by a model using only predictors in *S*. In all cases we used non-negative linear least-squares regression^83^ (N-fold cross-validated across condition pairs) to fit the lower triangle the relational dissimilarity matrices (RDMs) of models to RDMs of neural data. RDMs were computed as the euclidean distance between all pairs of conditions. For neural data, these RDMs were calculated in the high-dimensional neural space using the average firing rate of all neurons recorded within each brain area (pooled across sessions).

This formula evaluates the change in model performance, 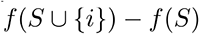, when predictor *i* is added to subset *S*, and weights each marginal contribution, ensuring equal representation of all subset sizes. Since Shapley values are calculated across a full set of subsets, each predictor’s contribution was normalized to the model’s total performance across all predictors. To match the amount of measurement noise between conditions, we took the most conservative approach and randomly downsampled all conditions to contain the same number of trials. We repeated the entire Shapley value calculation 10 times with different random subsamples of trials and averaged the result.

For significance testing, after trial averaging we randomly shuffled the conditions for each neuron independently and repeated the entire Shapley value calculation 100 times, using this distribution as a null distribution. A predictor’s Shapley value was considered significant if it exceeded the 99th percentile of this null distribution.

#### Motor control model

We trained a number of neural network models to control a biomechanical model of the arm by actuating simulated muscles during reaching using our previously developed open source toolbox, MotorNet^3^. For all models, the timestep size was 0.01 s, and we included a proprioceptive delay (20 ms), a visual delay (70 ms), and a muscle output delay (20 ms). We additionally included Gaussian noise at all time steps in the proprioceptive signal (SD: 1e-3), vision signal (SD: 1e-3), and in the muscle activation signal (SD: 1e-4).

Effectors were actuated using numerical integration with the Euler method. The arm26 model used in this study is available under the RigidTendonArm26 Effector class. It is briefly described below. The skeleton of the arm26 models follow the formalization proposed in Gomi and Kawato^84^, equations 1, 3, 5–7. The full formalization of the Hill-type muscles can be found in Thelen^85^, equations 1–7, and with the parameter values used in that study. When different parameters were provided for young and old subjects, the values for young subjects were used. In the RigidTendonArm26 class the moment arms are approximated as described in Kistemaker et al.^86^, equations A10–A12.

#### Recurrent neural network architecture

All networks consisted of one layer of GRUs with 256 units and standard activations (update/reset: sigmoid, candidate: tanh). Kernel and recurrent weights were initialized using Glorot initialization^87^ and orthogonal initialization^88^, respectively. At all time points we included gaussian noise in the candidate activation (before nonlinearity, SD: 1e-3). Biases were initialized at 0. Fifty percent of GRU units (equivalent results if 100%) were connected to the output layer of one node per muscle with a sigmoid activation function. The output layer’s kernel weights were initialized using Glorot normalization, and its bias was initialized at a constant value of −3. Because the output activation function is a sigmoid, this initial bias forces the output of the policy to be close to 0 at the start of initialization, ensuring a stable initialization state. Fifty percent of GRU units (equivalent results if 100%) received task-related and feedback inputs and these units were non-overlapping with units connected to the output layer. As task-related inputs, networks received a delayed vector (70 ms delay) of (x, y) cartesian coordinates for the start position and target position, target size, directional elbow perturbation probability (-1 to 1), and a binary cues indicating when the elbow probability cues was on, resulting in a 7-element input vector.

Networks also received delayed feedback (20 ms delay) from the environment consisting of proprioceptive signals containing muscle length and velocity for each muscle, the (x,y) position of the endpoint (70 ms delay), and a non-directional perturbation pulse (equal to one 20 ms after the perturbation, otherwise 0), resulting in a 15-element feedback vector.

#### Network training

Networks received interleaved training on a random reaching task and a probabilistic perturbation task. In the random reaching task, trials consisted of delayed reaches between random locations in the reachable workspace, where movement started after an unpredictable mechanical perturbation (random uniform -2 to 2 Nm perturbations shoulder and elbow) and no probability cues were given. Target size was randomized (0 - 10 cm diameter). In the probabilistic perturbation task details closely matched the monkey experiment. The start/end location was 60 degrees shoulder and 80 degrees elbow angle, targets were 1.2 cm diameter, and perturbations were -1 or 1 Nm elbow perturbations. In all training there was no background load and the randomized timing of cues was similar to the monkey experiment. Fifty percent of trials were catch trials (no perturbation) to prevent unwanted premature movements. Each training iteration consisted of a batch of 64 trials, each 3 second seconds long, and we used the Adam^89^ optimizer with a learning rate of 3e-3. Each task was trained for 2000 iterations.

Networks were optimized using a total loss that was a weighted sum of individual loss components, each addressing different aspects of the model’s performance:

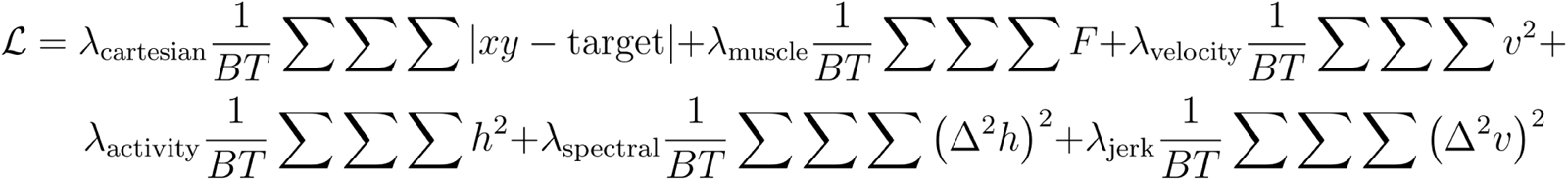

where *B* is the batch size, *T* is the total number of timesteps in an episode, *XY* and *target* are the current and target cartesian endpoints, *F* is the force applied by all muscles, *v* is the cartesian endpoint velocity, and *h* is the hidden activity of the network. Each component had a specific weight during training, specifically, 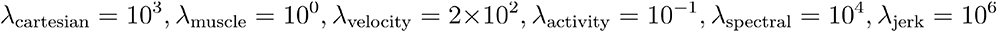 . The specific values of each of these components was not critical to successful training, with a few caveats. The muscle force penalty was necessary to prevent the network from simply using high levels of force at all times, the spectral penalty was necessary to prevent networks from learning chaotic dynamics as a results of the delayed sensory feedback, and the jerk penalty sped up training by encouraging networks to respond robustly to mechanical perturbations.

## Acknowledgements

Experiments were funded by: CIHR Operating Grants (Foundation Grant to J.A.P: 353197; Project Grant to J.A.P and J.D.: PJT-175010), Simons Foundation (via the Simons-Emory International Consortium on Motor Control), Azrieli Foundation (via the Collaboration on Motor Planning, Execution and Resilience). J.A.M. was supported by a Banting Postdoctoral Fellowship, a Vector Institute Postgraduate Affiliation, and a BrainsCAN Postdoctoral Fellowship (CFREF). J.A.P. was supported by the Canada Research Chair program.

## Author Contributions

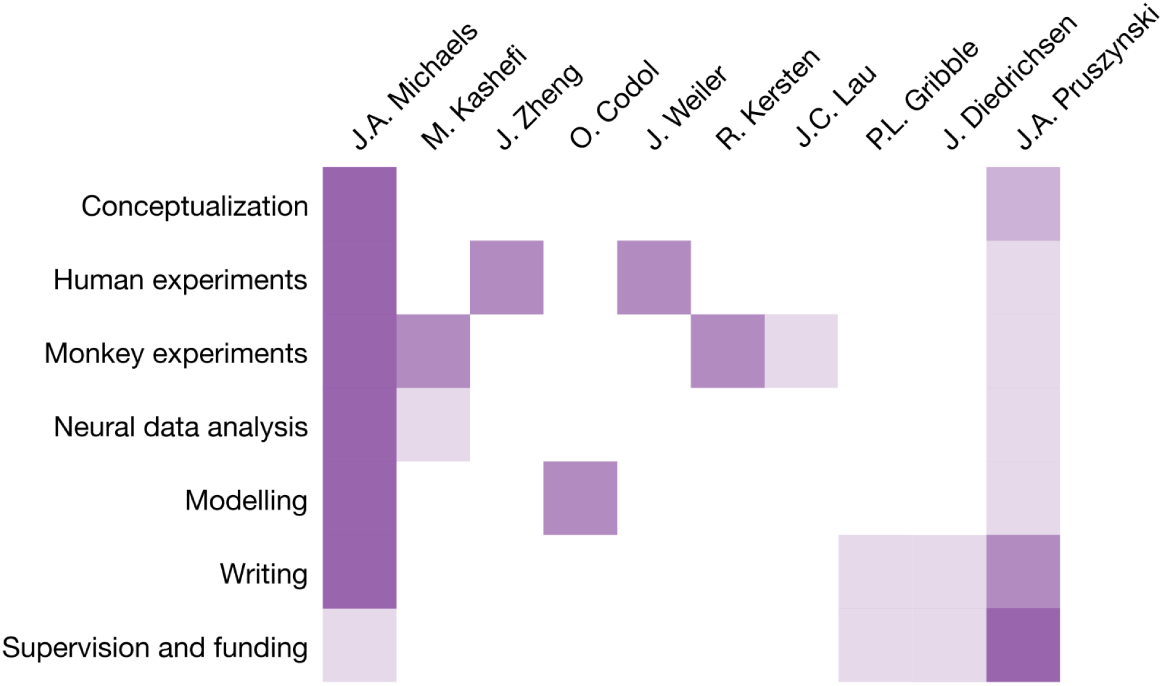

## Data availability

Data will be shared upon reasonable request by the corresponding author.

## Code availability

Custom codes for data analysis were written in MATLAB and Python and are available from the corresponding author upon request.

---

**Supplementary Fig. 1.**
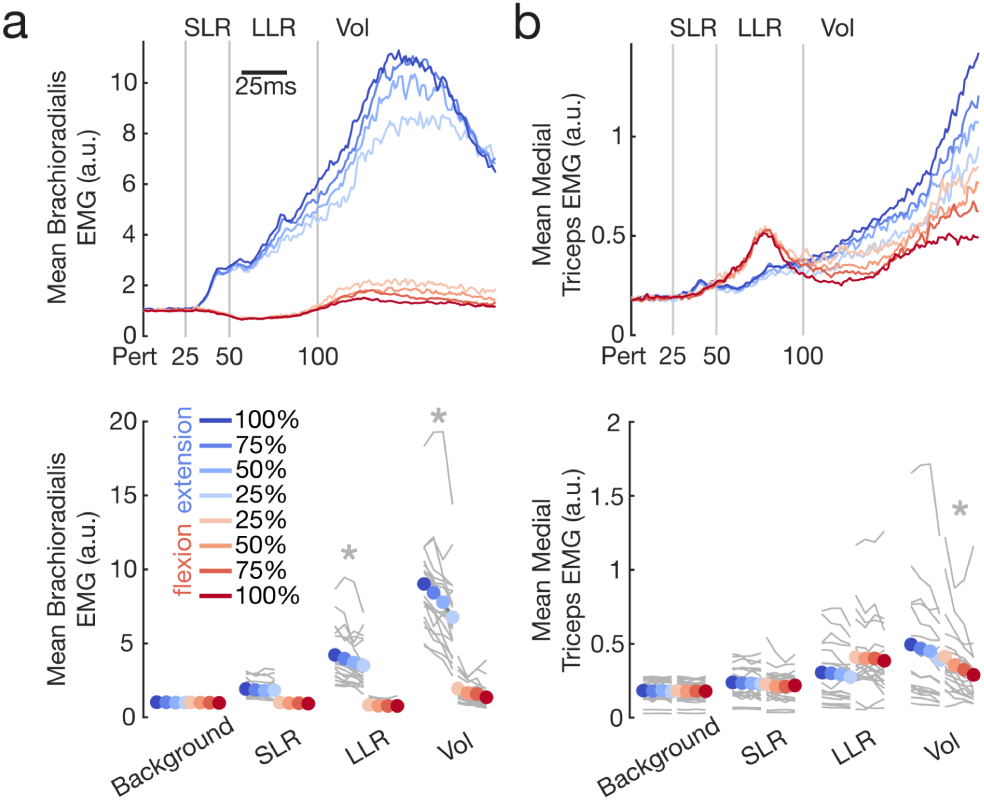
| Perturbation responses in human brachioradialis and medial triceps. a, In elbow extension conditions, no significant differences were found between probabilities in the background epoch (-200-0ms before perturbation) of the brachioradialis (F(3,19) = 1.57, p = 0.21) or in the short latency (SLR, 20-50ms) response (F(3,19) = 1.73, p = 0.17). In contrast, there was a significant effect of probability on EMG activity in the long latency epoch (LLR, 50-100ms) of brachioradialis (F(3,19) = 7.04, p = 0.0004) and during the voluntary epoch (100-150ms) of brachioradialis (F(3,19) = 32.62, p <0.0001). **b,** In elbow flexion conditions, no significant differences were found between probabilities in the background epoch (-200-0ms before perturbation) of the medial triceps (F(3,19) = 0.31, p = 0.82), in the short latency (SLR, 20-50ms) response (F(3,19) = 1.30, p = 0.28), or in the long latency epoch (LLR, 50-100ms) of the medial triceps (F(3,19) = 0.99, p = 0.40). In contrast, there was a significant effect of probability on EMG activity during the voluntary epoch (100-150ms) of the medial triceps (F(3,19) = 10.07, p < 0.0001).

**Supplementary Fig. 2.**
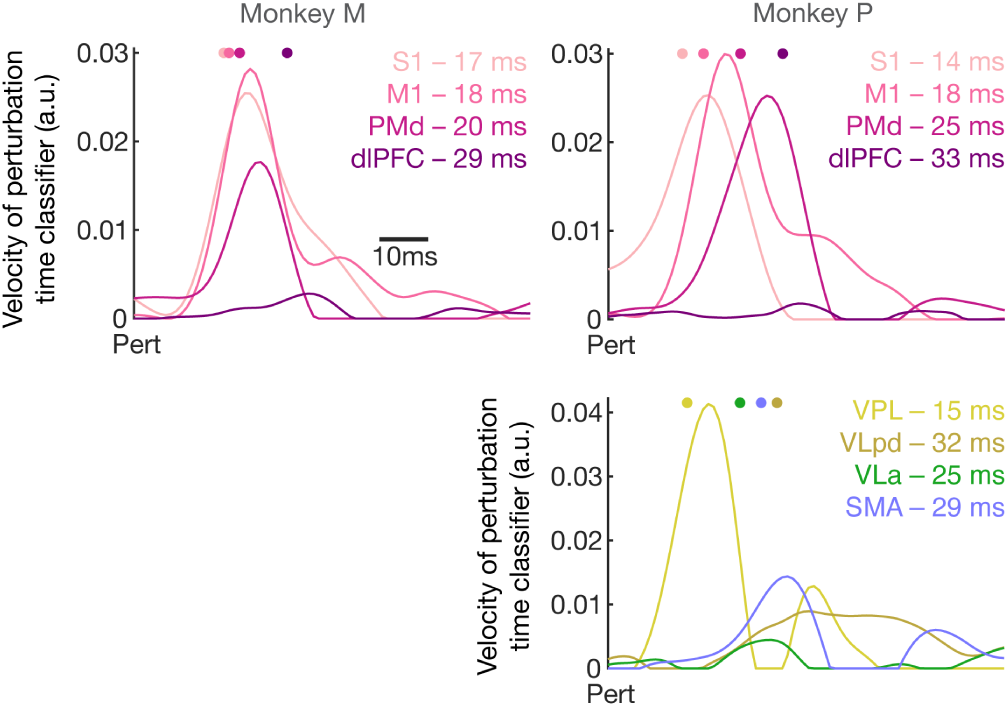
| Perturbation detection time across areas. For each brain area, we analyzed neural population data to determine the earliest time the perturbation could be detected. For each recording session, we fit a classifier to distinguish pre- and post-perturbation times across all trials within a session (SVM, 5-fold cross-validated, -150 to 150 ms relative to perturbation onset). We averaged the output of the classifier across all trials within each area, smoothed the result conservatively (50 Hz low-pass 4th-order zero-phase butterworth), and looked for positive peaks in the rates of change of the classifier output. The perturbation detection time was set at the point when velocity reached 80% of the max peak (colored dots above each plot).

**Supplementary Fig. 3.**
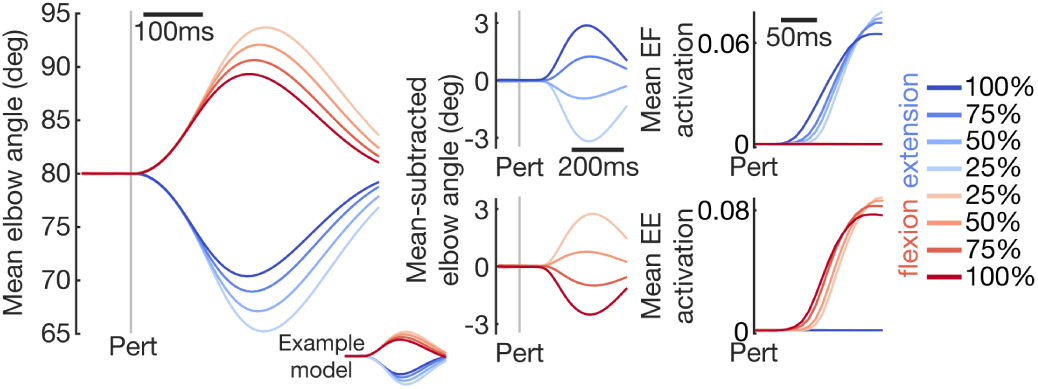
| Models with one-hot probability cue inputs develop sensory expectations. We trained 32 recurrent neural networks to control a biomechanical model of the arm during reaching, including realistic muscles, feedback, and delays, but using one-hot inputs to represent each probability cue (separate input channel for each cue) instead of the direct probability representation used in Figure 6. Average kinematics and muscle responses were virtually identical to the main results of Figure 6.

**Supplementary Fig. 4.**
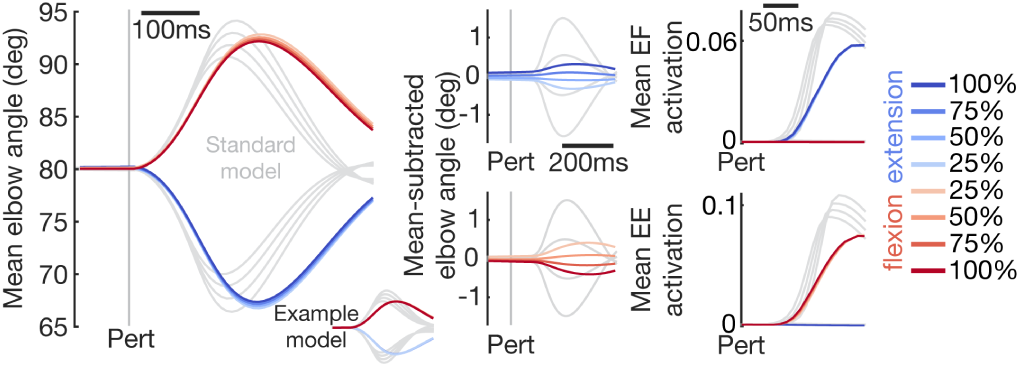
| Models without a condition-independent perturbation signal don’t express sensory expectations. We trained 32 recurrent neural networks to control a biomechanical model of the arm during reaching, including realistic muscles, feedback, and delays, but omitting the condition-independent perturbation pulse included in the results of Figure 6. Average kinematics of models performing the experiment did not scale with probability, nor did muscle activation within the long-latency reflex window (50-100 ms post-perturbation). The results of Figure 6 are outlined in gray.

